# Global signatures of Ribonucleotide Excision Repair deficiency and its therapeutic vulnerabilities

**DOI:** 10.64898/2026.10.09.757935

**Authors:** Ping Jing Toong, Soraya Boushaki, Salome J Zhao, Rachel Brough, Fei Fei Song, Dragomir B Krastev, Daniella Black, Giuseppe Rinaldi, Cherif Badja, Andrea Degasperi, Jan Czarnecki, Yasin Memari, Helen R Davies, Daniel Hübschmann, Hanno Glimm, Stefan Fröhling, Andrew N. J. Tutt, Stephen J Pettitt, Christopher J Lord, Gene C C Koh, Serena Nik-Zainal

## Abstract

Cancer therapies increasingly exploit DNA repair deficiencies, yet many clinically actionable repair-deficient states remain undetectable because they lack robust biomarkers and are not identifiable through gene-based diagnostics. Here, we show that deficiency of ribonucleotide excision repair (RER), a pathway that removes misincorporated ribonucleotides from genomic DNA, generates a distinct genome-wide mutational phenotype comprising characteristic insertion-deletion, substitution, and structural variation signatures. Using these features, we developed RERDetect, a whole-genome classifier that identifies RER-deficient tumors across diverse cancer types. Application of RERDetect to more than 11,000 primary and metastatic cancers revealed a recurrent subset of tumors with evidence of RER deficiency, the majority lacking detectable genetic alterations in *RNASEH2* genes. We identified promoter hypermethylation of *RNASEH2C* as a recurrent mechanism of pathway inactivation. RER-deficient cells exhibited distinct vulnerabilities, including sensitivity to *EXO1* and *FANCM* inhibition. These findings establish RER deficiency as a previously unrecognized and therapeutically actionable DNA repair-deficient state in cancer, demonstrating the power of mutational scars to identify clinically relevant tumor phenotypes that evade conventional genomic diagnostics.

## Main Text

Mutational scars provide a record of DNA damage and repair processes operating during tumor evolution and have become increasingly important biomarkers for cancer diagnosis and treatment (*1, 2*). In some cases, these scars identify clinically actionable repair-deficient states more effectively than gene-based diagnostics alone, revealing tumors that share common biological vulnerabilities despite diverse underlying mechanisms. However, many DNA repair and replication-stress defects lack well-defined genomic biomarkers, limiting their clinical recognition. This inability to detect cancers that carry specific DNA repair defects means that opportunities to therapeutically target cancers, for example, via DNA repair synthetic lethal effects, are often missed.

Ribonucleotide excision repair (RER), mediated by the heterotrimeric RNase H2 complex, removes ribonucleotides that are frequently misincorporated into genomic DNA during replication (*3, 4*). Failure of this pathway leads to persistence of embedded ribonucleotides and RNA:DNA hybrids, promoting genome instability (*5, 6*). In the absence of RNase H2, topoisomerase 1 (TOP1) processes embedded ribonucleotides to generate characteristic short 2-5bp deletions (*7, 8*), a mutational process previously linked to the cancer-associated indel signature COSMIC ID4, referred to as TOP1-dependent transcription-associated mutagenesis (TAM) (*9*).

Using a higher-resolution indel classification framework (*10*), we identified a distinct deletion signature InD4c, which occurs as a hypermutator phenotype in tumors and is accompanied by additional genome-wide mutational features. Here, we combined isogenic human cell models, pan-cancer whole-genome sequencing, and functional genetic screens to define the mutational consequences of RER deficiency (RERd). We show that RER deficiency generates a distinctive mutational phenotype that identifies a previously unrecognized, therapeutically actionable class of cancers, establish *RNASEH2C* promoter hypermethylation as a recurrent mechanism of pathway inactivation, and develop a genomic classifier for detecting this repair-deficient state from tumor genomes.

## Results

### RNASEH2-null cells produce global mutational signatures of all classes

To determine whether RER deficiency produces signatures across multiple variant classes, we generated isogenic *TP53*-null hTERT-immortalized RPE1 cell lines with CRISPR-mediated knockouts of each RNase H2 subunit: Δ*RNASEH2A*, Δ*RNASEH2B*, and Δ*RNASEH2C* (**Fig. 1A, fig. S1, A to D**). Validated knockout clones underwent 100-day mutation accumulation before 2–3 daughter subclones were isolated per genotype for whole-genome sequencing and mutational signature analysis **(Tables S1 and S2)**.

**Fig. 1.**
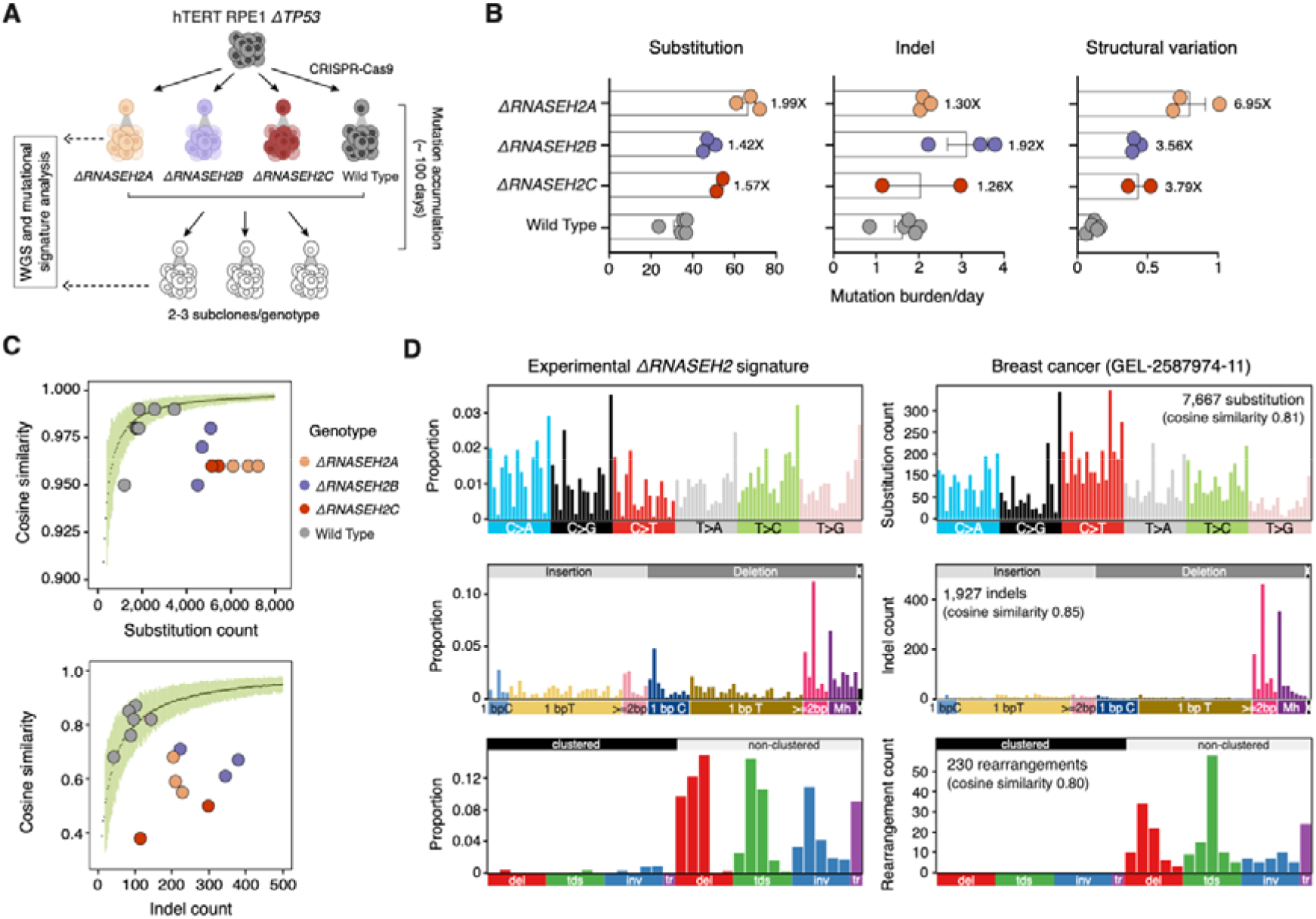
RNASEH2 knockouts induce distinct global mutational signatures. (A) Mutation accumulation experiment in *TP53*-null hTERT-immortalized retinal pigment epithelial cell (hTERT-RPE1 Δ*TP53*). *RNASEH2* knockout parental clones and an unedited wild-type clone (background) were cultured for 100 days. **(B)** Substitution, indel, and structural variation average fold increase in burden in edited subclones relative to wild-type subclones (*n* = 2-3 subclones per genotype; **Tables S1-3**). Mutation burdens were normalized by the number of days in culture. **(C)** Substitution and indel profiles of edited subclones are distinct from the background control. Light green error bars depict the mean ±3□s.d. of cosine similarities of *n* = 100 bootstrapped profiles generated from the average of *n* = 7 unedited wild-type subclones across a range of potential burdens. **(D)** Average background-subtracted experimental signatures of RPE1 Δ*RNASEH2A*, Δ*RNASEH2B*, and Δ*RNASEH2C* (*n* = 8 subclones) on left, and mutational catalogues from a breast cancer GEL-2587974-11 sample with high InD4c exposure on right.

RNASEH2 loss increased all major classes of mutations, including single-base substitutions, indels, and structural variants, compared to wild-type controls, indicating that the consequences of RER loss are not confined to deletional mutagenesis (**Fig. 1B** and **Table S3**). The mutational profiles of Δ*RNASEH2A*, Δ*RNASEH2B*, and Δ*RNASEH2C* subclones were highly concordant with one another and clearly distinct from unedited controls (**Fig. 1C**), supporting a shared pathway-level phenotype rather than a subunit-specific effect.

Across the three knockout genotypes, RER deficiency was defined by a composite genome-wide mutational pattern encompassing: i) an SBS3-like substitution signature (cosine similarity 0.84), ii) an indel signature characterized by deletions of 2-4 bp at non-repeats, deletions of 2-8 bp at short tandem repeats (2-4 repeat units), and deletions between 2–5 bp with 1 bp of microhomology, most similar to cancer-derived InD4c (cosine similarity 0.83), and iii) a distinct structural variation signature comprising long (100 kb–1 Mb) deletions, tandem duplications, and inversions, resembling the rearrangement features of InD4c-high tumors (**Fig. 1D, fig. S1E**, and **Tables S4-S6**). Thus, RNase H2 loss leaves a distinct genome-wide mutational scar that extends beyond the short deletion phenotype previously associated with defective ribonucleotide processing (*7-9*).

Because this mutational phenotype included an SBS3-like pattern and elevated microhomology-mediated deletions, both widely used markers of HRD, we evaluated HR activity in cells with RNase H2 loss.

### SBS3 in RNASEH2-null cells is not due to HR deficiency

We measured HR directly in matched isogenic Δ*RNASEH2A*, Δ*RNASEH2B*, and Δ*RNASEH2C* cells using an extrachromosomal reporter assay (*11*). All three knockouts retained HR activity indistinguishable from wild-type cells, in stark contrast to the profound HR impairment observed in Δ*BRCA1* cells generated in the same genetic background (*12*) (**fig. S2A**). This finding is consistent with prior work showing that RNase H2-deficient cells maintain RAD51 foci formation and near-wild-type DR-GFP activity despite marked sensitivity to PARP inhibition (*13*). The PARP inhibitor sensitivity of RNase H2-deficient cells, which was recapitulated in our *RNASEH2* knockout models (**fig. S2B**), is thought to not arise from an HR defect but from persistent genomic ribonucleotides (increased in RNase H2-deficient cells) ultimately being processed into PARP1-binding DNA substrates. In the presence of PARP inhibitor, PARP1 is “trapped” at these substrates, causing cytotoxicity (*13*).

Global transcriptomic profiling further distinguished RNase H2 loss from canonical HR deficiency (HRD). Relative to Δ*BRCA1* cells, Δ*RNASEH2* cells showed a distinct transcriptional response, including upregulation of DNA recombination pathways, consistent with compensatory or hyper-recombinogenic activity rather than recombination failure (**fig. S2, C and D**). Together with previous observations of increased sister chromatid exchange in RNase H2-deficient cells and the synthetic lethality between *RNASEH2B* and *BRCA1/BRCA2* (*13*), these data indicate that RER deficiency does not phenocopy *BRCA1* loss at the level of HR function.

Accordingly, the SBS3-like signature in RER-deficient cells cannot be attributed to impaired HR. Although Δ*RNASEH2* and isogenic Δ*BRCA1* or Δ*BRCA2* cells showed comparable SBS3 exposures, their indel spectra diverged markedly: RERd genotypes were enriched for deletions with 1-bp microhomology, characteristic of InD4c, whereas *BRCA1-* or *BRCA2*-deficient cells showed the longer microhomology-associated deletions typical of InD6 (*10, 14*), with a mode of ≥2 bp microhomology at the indel junction (**fig. S2E, fig. S3A**). These findings highlight how SBS3 is not specific to HR deficiency and can also arise in HR-proficient RER-deficient cells.

### RER-deficiency InD4c is distinct from COSMIC ID4 and generates a hypermutator phenotype

We next examined the relationship between the indel pattern derived from *RNASEH2*-null cells and the published cancer-derived indel signatures linked to TOP1-associated mutagenesis, including COSMIC ID4, MuSiCal ID4, and InD4a (*9, 10, 15*). We previously found that our 89-channel indel classification system (*10*), which has a higher-resolution than the COSMIC 83-channel system, could decompose InD4a into two patterns: InD4c, present as a hypermutator phenotype in tumors, and InD4d (a residual component of InD4a) (**fig. S4A)**. COSMIC ID4 more closely matched mutation profiles of cancers enriched with InD4a, whereas MuSiCal ID4 matched InD4c-enriched tumors (**fig. S4B, Table S7**). Crucially, the experimentally derived Δ*RNASEH2* indel profile was more similar to InD4c and to MuSiCal ID4 (cosine similarities 0.93 and 0.86, respectively) than to COSMIC ID4 (cosine similarity 0.82) (**fig. S4C**). Together, these data indicate that the RER-deficient indel phenotype is captured more specifically by InD4c (and by MuSiCal ID4) than by COSMIC ID4.

We then asked whether InD4c-high tumors were biologically distinct from tumors dominated by InD4a/COSMIC ID4. We examined the sequence motif of 2-bp deletions, which is a prominent feature shared by both signatures. Similar to cancers with InD4a exposures, 2-bp deletions in InD4c cancers were enriched at the TOP1-associated TNT sequence motif (deleted bases underlined) and in transcribed regions, consistent with a shared link to TOP1-dependent transcription-associated mutagenesis (**fig. S5**) (*9*). However, despite these shared features, the composition of 2-bp deletions within the TNT motif differed between the two groups: InD4c-high tumors showed a higher proportion of CT deletions than InD4a-high tumors (**fig. S5B)**, which may be explained by the preferential incorporation of ribouridine at CT/GT dinucleotides (*9, 16*). The lack of CT enrichment in InD4a cancers therefore suggests that InD4a/COSMIC ID4 may be associated with increased TOP1 activity during increased transcription rather than from pathologically incurred increased ribonucleotide incorporation caused by RER deficiency. Experimental data recapitulated the cancer data (**fig. S5B**), reinforcing this postulate.

Furthermore, comparing the prevalence of InD4a and InD4c across 12,007 high-depth whole-genome-sequenced primary cancers from the Genomics England (GEL) cohort, we found that InD4a was common but typically low-level, being detectable in 3,827 tumors (31.9%), whereas InD4c was present in only 126 tumors (1.05%) but at substantially higher exposure (*P*<0.0001, **fig. S4D**). Only 1.75% of tumors with detectable InD4a exposures had more than 800 indel mutations assigned to the signature, compared to 15.1% of tumors with InD4c exposures. InD4c tumors also carried significantly higher SBS3 exposures (*P*<0.0001) and structural variant burden (*P*<0.0001) compared to InD4a tumors (**fig. S4D**). Thus, although InD4a and InD4c both have features of TOP1-associated deletional mutagenesis, InD4a likely reflects a generic, low-level TOP1/transcription-associated deletion process, whereas InD4c, recapitulated by isogenic Δ*RNASEH2* cells, represents a distinct subtype associated with pathological RER deficiency.

### RER deficiency produces diverse, predominantly long structural variants

RNASEH2-null cells also showed diverse structural variants, including deletions, tandem duplications, and inversions, mostly occurring at larger sizes (100kb-1Mb), together with some translocations (**Fig. 1D**). This pattern was distinct from the rearrangement profiles of isogenic Δ*BRCA1* cells, in which tandem duplications and deletions were markedly shorter, typically below 10 kb (**fig. S3A**). Thus, although RER deficiency shared substitution features with HRD, its SV phenotype was qualitatively different. This observation was reinforced in InD4c-high cancers, which showed an approximately equal distribution of all SV classes, with tandem duplications, deletions, inversions, and translocations at roughly a 1:1 ratio – in marked contrast to *BRCA1*-mutant cancers, where tandem duplications outnumbered deletions by twofold and exceeded inversions by nearly fivefold (*P* < 0.0001) (**fig. S6A**).

Long tandem duplications were not unique to RER deficiency; they were also observed in *CDK12*-mutant and *CCNE1*-amplified tumors (**fig. S6B**). However, these other genetic contexts lacked the full spectrum of deletions, inversions, and translocations seen in RERd, and did not show the accompanying InD4c and SBS3 features that characterize it.

The overall experimentally derived rearrangement pattern had only modest similarity to any single cancer-derived rearrangement signature (cosine similarities < 0.78). Instead, the rearrangement pattern comprised several components that individually resemble features from some rearrangement signatures, including the short deletion signature R5 (0.77), long tandem duplication signatures R1 (0.52) and R14 (0.34), and long deletion signature R7 (0.72) (**fig. S6, C and D**). This wide distribution across multiple rearrangement signatures suggests that failure to remove misincorporated ribonucleotides by RER may lead to a generic, initiating DNA damage lesion, capable of producing multiple SV classes.

To test this hypothesis more directly, we examined whether InD4c deletions were spatially associated with known hotspots of long tandem duplication formation. Thirty-three genomic hotspots for long (>100 kb) tandem duplications have previously been described (*17*), attributed to a high propensity for DNA damage at sites of elevated transcription. Because ribonucleotide misincorporation is directly linked to transcription-associated mutagenesis, we asked whether InD4c deletions were enriched at these same loci. We found a significant enrichment of InD4c deletions at the 33 previously described tandem duplication hotspots (*P* < 0.0001), and an agnostic hotspot analysis independently identified 253 InD4c hotspots, of which 16 overlapped with 27% (9/33) of these sites (*P* < 0.0001; **fig. S6E**). This spatial convergence supports a mechanistic link between InD4c deletions and SV formation at transcriptionally active genomic regions.

Finally, we examined the repair mechanisms operating at SV breakpoints in RER-deficient versus HR-deficient cells. The distribution of microhomology lengths at tandem duplication breakpoints differed markedly between Δ*RNASEH2* cells and both Δ*BRCA1* cells and *BRCA1*-mutant cancers (*P* < 0.0001; **fig. S6F**). *BRCA1*-deficient contexts showed a characteristic peak of ∼2-bp microhomology at tandem duplication breakpoints, consistent with microhomology-mediated end joining as the dominant repair mechanism underlying rearrangement signature R3 (*2*). This peak was absent in Δ*RNASEH2* cells, suggesting a different balance of compensatory end-joining mechanisms at tandem duplication breakpoints in RER deficiency and further distinguishing the SV landscape of RERd from that of classical HRD.

### RERDetect: a genomic classifier for RER deficiency

The composite RERd mutational phenotype encompassing SBS3, InD4c, and a distinct SV signature overlaps substantially with that of HRD, creating a dual problem: existing classifiers (*18, 19*) such as HRDetect (*20*) and HRProfiler (*21*) misclassify RER-deficient tumors as HR-deficient, while some RER-deficient tumors remain undetected in the absence of a dedicated classifier (**fig. S7**). Indeed, among HRDetect-high and -intermediate cancers lacking canonical HR gene mutations, InD4c-positive cases were significantly enriched (*P* = 0.0001; **fig. S7, A to D**). To address this gap, we developed RERDetect, a lasso logistic regression classifier trained on the experimentally defined mutational features of Δ*RNASEH2* cells described above.

The training dataset comprised 19 experimental Δ*RNASEH2* samples as positives and 208 samples lacking *RNASEH2* mutations as negatives, spanning diverse DNA repair mutants and wild-type genotypes (**Fig. 2A**). We trained and compared three classes of models using different SNV, indel, and rearrangement features as inputs: i) mutational catalogue contexts, ii) absolute signature exposures, and iii) signature exposures as proportions. All models were trained using lasso logistic regression with 10-fold cross-validation, repeated across 10 independent 70:30 random partitions to assess feature stability, then retrained on all 227 samples (**Table S8**). The model using mutational catalogue contexts achieved the best discrimination, with a cross-validated AUROC (area under the receiver operating characteristic) of 1.0 (**Fig. 2B, Table S9**) and was selected as the final RERDetect model. Its seven discriminatory features, selected by lasso regularization, comprised four indel channels characteristic of InD4c (2-bp deletions at 1-bp microhomology and at 2-, 3-, and 4-unit short tandem repeats), two substitution channels (C[T>A]A and C[T>A]G), and 10-100 kb tandem duplications (**Fig. 2C, Table S10**), directly reflecting the multi-class nature of the RERd phenotype.

**Fig. 2.**
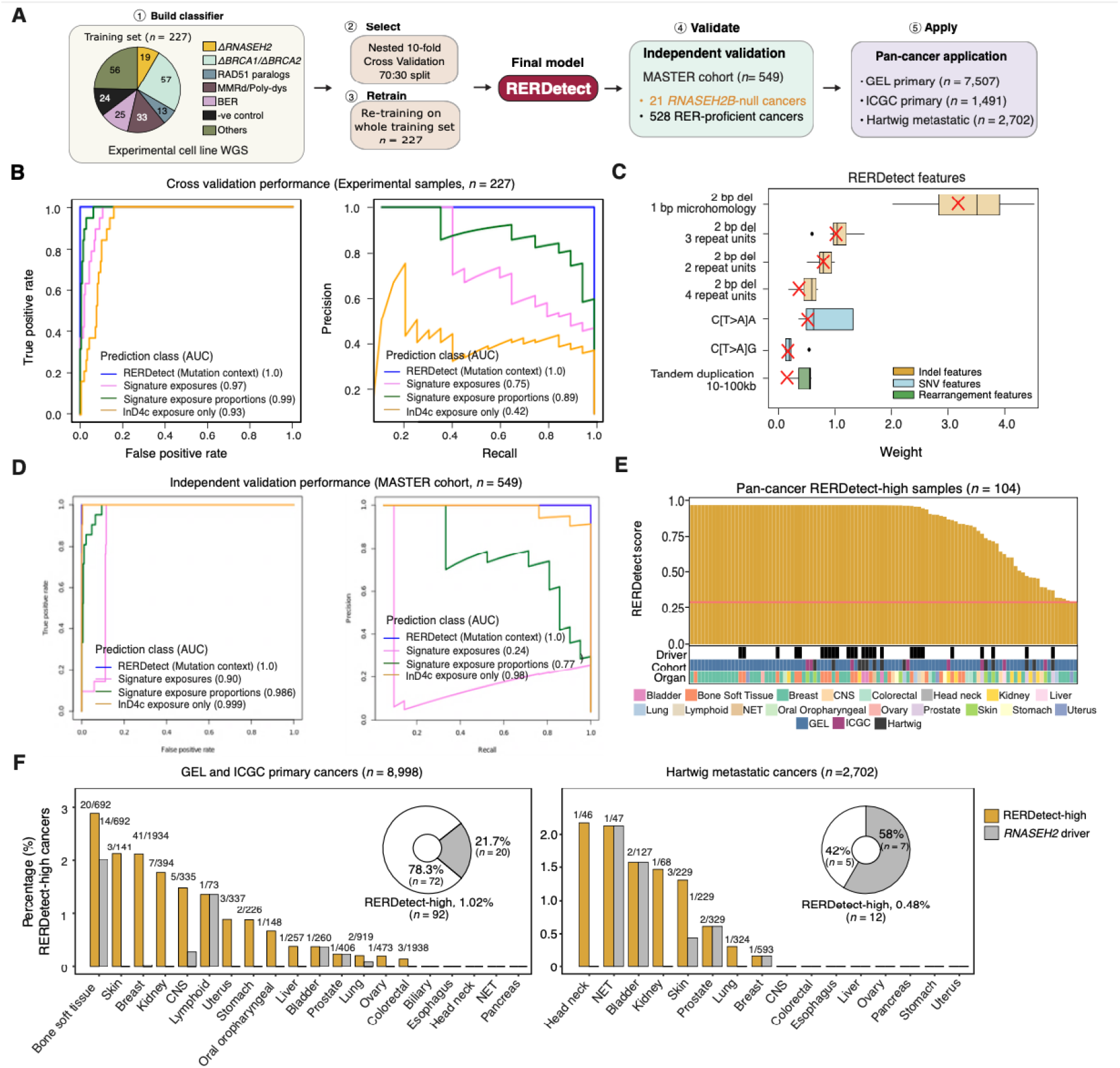
RERDetect: a genomic classifier for RER deficiency. (A) Simplified workflow for RERDetect development. (1) Initial training using logistic regression models with 227 different cell lines of diverse genotypes (2) Ten-fold cross validation with 70:30 partitioning to assess and select features. (3) Final retraining using all the samples to produce the best performing RERDetect classifier. (4) Models were validated on an independent cohort (*n* = 549). (5) Final model was applied to a large pan-cancer cohort of GEL (*n* = 7,507) and ICGC (*n* = 1,491) primary cancers and Hartwig metastatic cancers (*n*= 2,702). **(B)** Performance of models trained using different input features in the initial cross validation from experimental samples. **(C)** Distribution of coefficients for the seven genomic features contributing to the final RERDetect model. Boxplots depict the ten replicates of training in cross-validation. The red cross represents the final weights from the final retraining contributing to RERDetect. **(D)** Performance of the same models in independent validation MASTER cohort (*22*) (*n* = 549). **(E)** RERDetect-high cancers (*n* = 104) from GEL, ICGC, and Hartwig. Each vertical line represents one sample ordered from left to right by decreasing RERDetect score. The presence of an *RNASEH2* driver event and the cancer tissue origin are indicated below. **(F)** The percentage of cancers that are RERDetect-high (yellow) and have a driver mutation in any *RNASEH2* gene (grey) for each tissue type in primary (*n* = 8,998) and metastatic (*n* = 2,702) cancers.

We then evaluated all three models in an independent cancer cohort (*22*) comprising 21 samples with confirmed homozygous *RNASEH2B* deletions as positives and 528 *RNASEH2*-wild-type controls as negatives, selected to have no mutations in any *RNASEH2* gene and above average expression of all three *RNASEH2* subunits by RNA-seq. The concordance between predictions and annotations was assessed by calculating the AUROC and the area under precision-recall (AUPRC) curves. The RERDetect model based on mutation contexts achieved an AUROC and an AUPRC of 1.0 (**Fig. 2D, Table S11**), outperforming other models using signature exposure proportions (AUPRC 0.771) and InD4c exposure alone (AUPRC 0.984). The superiority over InD4c exposure alone demonstrates that integrating substitution and rearrangement features alongside the indel signals captures RERd more completely than any single signature. RERDetect also achieved a maximum F1-score (1.0) for predicting RER deficiency at a cutoff of 0.3 in this validation cohort; this cutoff was therefore set as the classification threshold.

We next applied RERDetect to high quality samples from three large pan-cancer WGS cohorts: 7,507 primary cancers from GEL, 1,491 from ICGC, and 2,702 metastatic cancers from Hartwig. Across 11,700 cancers, 104 (0.89%) were classified as RERDetect-high (**Fig. 2E, Table S12**), spanning 17 tissue types and most frequently occurring in cancers of bone and soft tissue, skin, breast, kidney, and CNS (**Fig. 2F)**. RERd prevalence was slightly lower in the metastatic cohort (0.44%) than in primary cohorts (GEL:1.12%; ICGC:0.54%), likely reflecting differences in cohort composition or treatment selection. Strikingly, only 27 of 104 cancers (26%) harbored a confirmed genetic driver event in *RNASEH2*. This finding suggests that most RER-deficient cancers cannot be explained by coding disruption of *RNASEH2* genes alone, raising the question of whether alternative mechanisms causing RER defects are operative in human cancer.

### RNASEH2C promoter hypermethylation underpins a subset of RER-deficient cancers

To characterize the landscape of driver events underpinning RERd, we analyzed the somatic genetic drivers of *RNASEH2* in the 104 RERDetect-high tumors (**Fig. 3A, Table S13**). Only 26 (25%) had *RNASEH2B* deletions, predominantly in sarcomas, bladder cancer, prostate cancer, and chronic lymphocytic leukemia (CLL). *RNASEH2B* resides within the 13q14 locus – a region recurrently deleted in multiple cancer types and harboring several established tumor suppressors. Consistent with this, we noted 77% (20/26) of tumors with *RNASEH2B* deletion had co-loss of *DLEU1/2* along with the microRNA cluster miR-15a/16, *SETDB2*, and *RB1* (**Fig. 3, B and C**). This could be important for two reasons. First, miR-15a/16 has been implicated in regulating expression of *PD-L1, PD-1*, and *CTLA-4*, thus potentially this subset of RERd tumors could exhibit immune evasion (*23, 24*). Second, co-loss of *RB1* has implications for PARP inhibitor resistance (*25*). Regardless, all 26 cases had RERd mutational features, reinforcing the penetrance of mutational phenotype associated with *RNASEH2B* mutations. We also identified one breast cancer (GEL-2622518-11) with a germline deep deletion of *RNASEH2A* in this cohort. Unexpectedly, most primary tumors with confirmed *RNASEH2B* driver events had significantly lower InD4c burdens than driver-negative cases (mean exposure 285.4 vs. 588.1, *P* = 0.01357) (**Fig. 3D**), and InD4c hypermutators (burden >500) – comprising 27% of RERDetect-high primary cancers – rarely harbored genetic driver events. These observations indicate that canonical *RNASEH2* genetic inactivation accounts for only a minority of RER-deficient tumors and suggest that alternative drivers/mechanisms may be responsible for impairing RER function.

**Fig. 3.**
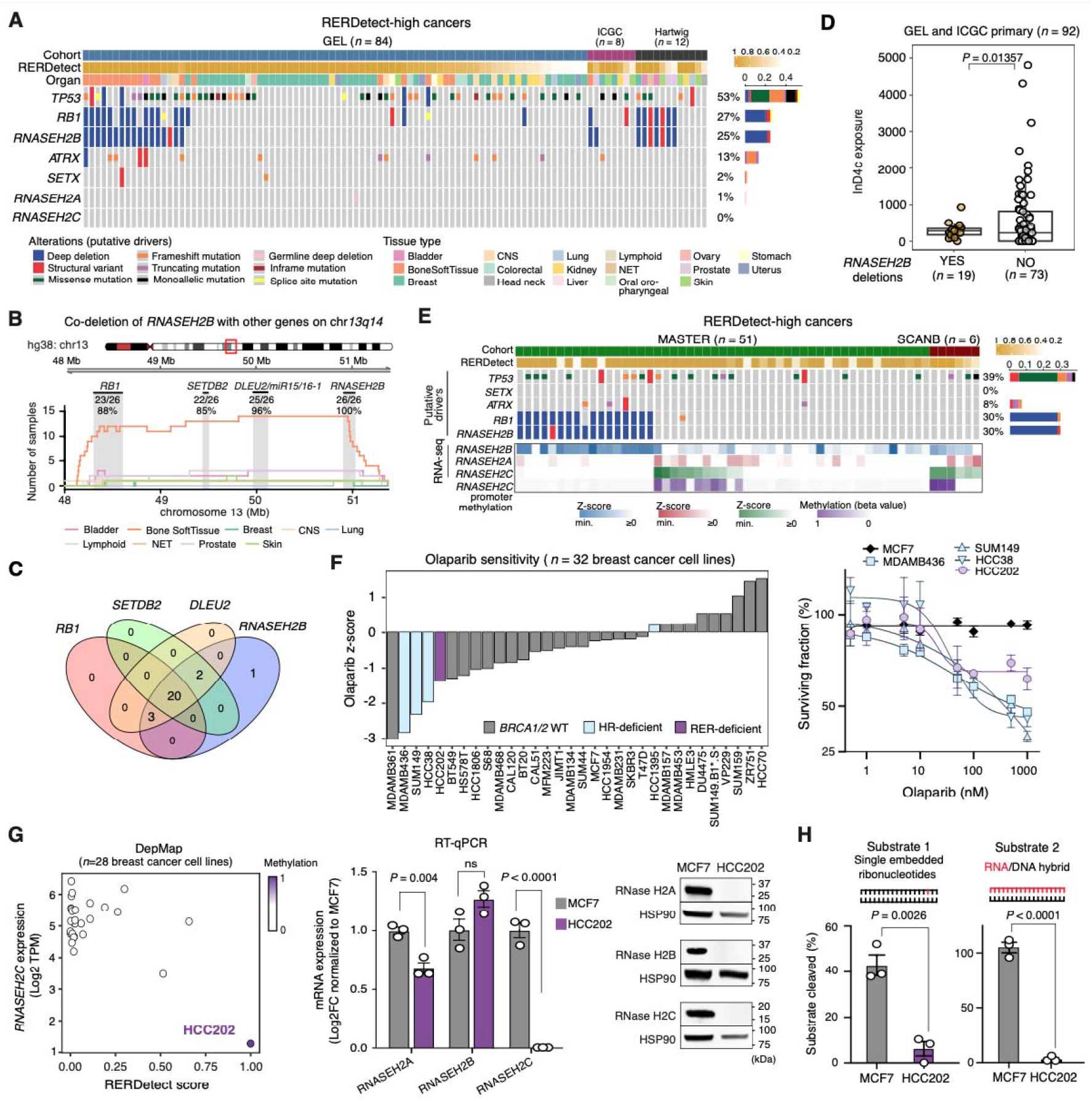
RNASEH2C promoter hypermethylation is a recurrent mechanism of RER deficiency. (A) Oncoprint of RERDetect-high cancers (*n* =104) in GEL, ICGC, and Hartwig showing frequency of driver mutations in *RNASEH2* genes, *TP53*, and helicases *ATRX* and *SETX*. **(B)** Deletion density plot showing number of RERDetect-high cancers with homozygous deletions spanning the 48-52 Mb region on chromosome 13. **(C)** Venn diagram showing number of RERDetect-high cancers with co-loss of *RNASEH2B, DLEU1/2, miR15a/16-1, SETDB2*, and *RB1* on chromosome *13q14*. **(D)** The InD4c exposures of RERDetect-high samples in primary cancers (GEL/ICGC) with (*n* = 19) and without (*n* = 73) homozygous *RNASEH2B* deletion. **(E)** Oncoprint of RERDetect-high cancers in the MASTER (*n* = 51) and the SCAN-B (*n* = 6) cohorts showing the frequency of genetic driver mutations, the gene expression of *RNASEH2* genes, and the methylation levels at the *RNASEH2C* promoter. The RNA-seq gene expression is z-score normalized log2 TPM. **(F)** Waterfall plot of olaparib AUC Z-scores (left) and dose-response SF_50_ curves (right) of olaparib across breast cancer cell lines (*28*). **(G)** *RNASEH2C* gene expression and RERDetect score of *n* = 28 DepMap breast cancer cell lines (left) (*27*). *RNASEH2C* gene expression by qPCR was significantly decreased in HCC202 compared to MCF7 (*P* < 0.0001; multiple unpaired t-tests with Benjamini Hochberg correction; middle). Immunoblotting of whole-cell extracts from MCF7 and HCC202 for RNase H2A, RNase H2B, and RNase H2C (right). **(H)** HCC202 showed significantly decreased enzymatic activity towards substrate 1 (dsDNA with single embedded ribonucleotide, *P* = 0.0026) and substrate 2 (RNA-DNA hybrids, *P* < 0.0001). *P* values were calculated using an unpaired two-tailed t-test.

Searching for recurrent alterations in other cancer genes, including *TP53* and helicase genes, across RERDetect-high cases yielded no strongly recurrent candidates (**Fig. 3A**). We therefore asked whether epigenetic mechanisms could drive the InD4c phenotype. Because the GEL cohort does not have methylation data, we analyzed the MASTER cohort of rare cancers (*22*) and the SCAN-B cohort of triple-negative breast cancers (*26*), both of which have matched WGS, transcriptomic, and methylation data. In the MASTER and SCAN-B cohorts, respectively, we identified 9 of 51 (17.6%) and 3 of 6 (50%) RERDetect-high cancers with significantly high promoter hypermethylation scores across the *RNASEH2C* CpG island, which correlated with concordantly low *RNASEH2C* gene expression (**Fig. 3E**). All 12 cases exhibited clear InD4c, long tandem duplications, and SBS3 exposures, with no other *RNASEH2* genetic driver, nominating *RNASEH2C* promoter hypermethylation as a candidate epigenetic driver of RERd in human cancer.

To validate this finding, we identified HCC202, a breast cancer cell line within the DepMap project (*27*), as RERDetect-high (0.99), exhibiting all hallmark signatures of RERd. Re-analysis of published PARPi sensitivity data involving 32 breast tumor cell lines (*28*) showed HCC202 to be as olaparib-sensitive as *BRCA1*-mutant lines (MDAMB436, SUM149) and the *BRCA1* promoter-hypermethylated HCC38 – despite harboring no detectable mutations in *BRCA1/BRCA2* or any *RNASEH2* genes (**Fig. 3F**). Publicly available methylation and transcriptomic data (*29*) demonstrated near 100% CpG methylation at *RNASEH2C* promoter and profound downregulation of *RNASEH2C* expression in HCC202 compared to other breast cancer cell lines, which we independently validated using bisulfite-specific PCR and RT-qPCR (**Fig. 3G**). Interestingly, although only *RNASEH2C* was transcriptionally repressed in HCC202, we observed near-total loss of all three RNase H2 subunits at the protein level, consistent with destabilization of the obligate RNase H2 heterotrimer upon loss of a single subunit (**Fig. 3G**). Furthermore, HCC202 lysates showed significantly reduced enzymatic activity against single embedded ribonucleotides (*P* = 0.0026) and RNA:DNA hybrids (*P* < 0.0001) (**Fig. 3H**). Together, these data implicate *RNASEH2C* promoter hypermethylation as a recurrent and previously unrecognized epigenetic driver of a subset of RER-deficient tumors, although additional mechanisms likely account for the remaining driver-negative cases.

### RERDetect predicts synthetic lethal effects distinct from classical HRD

Although *RNASEH2* defects have been associated with synthetic lethal effects also seen in classical HRD, namely PARPi sensitivity (confirmed in our Δ*RNASEH2A* cells, **fig. S2B**), the distinct mutational and functional profile of RERDetect-high cancers prompted us to assess whether RERd elicits a distinct set of targetable genetic dependencies. This could, in principle, enhance precision medicine intervention for both HRD and RERd cancers.

We thus carried out a genome-wide CRISPR dropout screen in Δ*RNASEH2A* cells, contrasting the results against six isogenic HRD-associated knockouts (*12*) (**Fig. 4, A and B; Table S14**). This parallelized approach allowed us to identify shared and unique synthetic lethal effects. Dependencies on *PARP1, ATR*, and *TOP1* can be caused either by HRD, or in *RNASEH2* defective cells, by impaired ribonucleotide excision repair (*13, 30*). Δ*BRCA1*, Δ*BRCA2*, and Δ*RNASEH2A* cells all shared synthetic lethality with loss of *PARP1, ATR*, and *TOP1*. Δ*RNASEH2A* cells also shared *FANCM* dependency with Δ*BRCA1* cells only (**Fig. 4, C and D**). Conversely, and consistent with the absence of HRD in Δ*RNASEH2A* cells, two HRD-dependent synthetic lethal effects involving *POLQ* (*31, 32*) and *CIP2A* (*33*) were seen in Δ*BRCA1*, Δ*BRCA2*, Δ*PALB2*, Δ*RAD51C*, and Δ*RAD51D*, but absent in Δ*RNASEH2A* cells (**Fig. 4, C and D**), suggesting that POLQ (*34, 35*) or CIP2A inhibition, unlike in classical HRD, might not be a promising therapeutic strategy in RER-deficient cancers. In *BRCA1-* or *BRCA2*-mutant cancers, increased usage of POLQ-mediated end joining as a favored mechanism of double-strand break repair correlates with enrichment of longer microhomology-associated deletions or InD6 (see earlier). The relative lack of InD6 in Δ*RNASEH2A* cells was therefore in keeping with the lack of *POLQ* dependency in these cells.

**Fig. 4.**
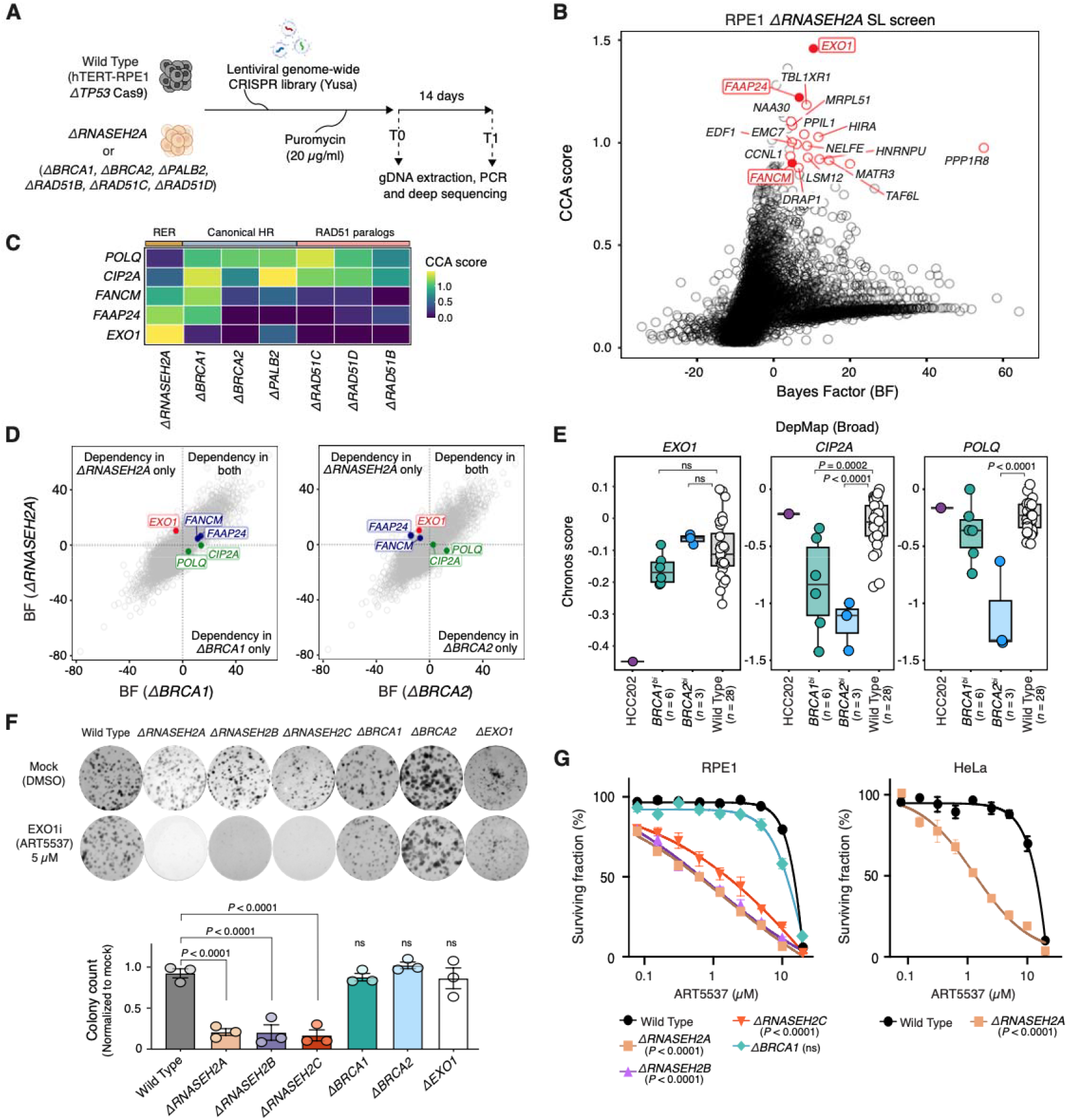
Synthetic lethality and therapeutic response in RERd. (A) Schematic of parallel genome-wide CRISPR dropout screens in hTERT-RPE1 Δ*TP53* (control) and isogenic Δ*RNASEH2A* to detect synthetic lethal interactions. **(B)** Scatterplot of CRISPR count analysis (CCA) and Bayes Factor (BF) derived from BAGEL2 for the Δ*RNASEH2A* clone. **(C)** Heatmap of CCA values of selected synthetic lethal genes in Δ*RNASEH2A* clones compared to isogenic HR gene knockouts (*12*). **(D)** BF scores of gene targets in Δ*RNASEH2A* clone against those of Δ*BRCA1* and Δ*BRCA2* clones. Some genetic dependencies were found only in Δ*RNASEH2A* (red), in both Δ*RNASEH2A* and Δ*BRCA1* (blue), or in only Δ*BRCA1* and Δ*BRCA2* (green). **(E)** *EXO1, CIP2A*, and *POLQ* dependency (Chronos score) in HCC202, *BRCA1*-mutant cell lines (*n* = 6), *BRCA2*-mutant cell lines (*n* = 3), and wild-type cell lines (*n* = 28) from the Broad DepMap screens (*27*). *P* values were calculated using one-way ANOVA with Tukey’s multiple comparisons test. **(F)** Clonogenic survival of hTERT-RPE1 Δ*TP53* (wild-type) and isogenic knockouts treated with EXO1 inhibitor (5 μM) or mock (DMSO control) for 7 days. The bars indicate the mean ±□s.d. (*n* = 3 replicates). Δ*RNASEH2A* (*P* < 0.0001), Δ*RNASEH2B* (*P* < 0.0001), Δ*RNASEH2C* (*P* < 0.0001). *P* values were calculated using one-way ANOVA with Dunnett’s multiple comparisons test. **(G)** Dose-response SF_50_ curves of EXO1 inhibitor ART5537 in hTERT-RPE1 Δ*TP53* wild-type and isogenic Δ*RNASEH2* clones and in HeLa wild-type and Δ*RNASEH2A* cell lines. Error bars represent mean ± s.e.m. (*n* = 3 replicates). *P* values were calculated using extra sum-of-squares F test of the IC_50_ with Bonferroni multiple comparisons correction.

Among synthetic lethal effects that were relatively unique to Δ*RNASEH2A* cells, the strongest candidates included the 5’ to 3’ exonuclease EXO1 and the Fanconi anemia pathway heterodimer, FANCM–FAAP24 (C19orf40). *EXO1* synthetic lethality was seen in Δ*RNASEH2A* cells but absent in the other isogenic HR gene knockouts (**Fig. 4, C and D**). Importantly, we found in DepMap data that HCC202 – the breast cancer cell line with *RNASEH2C* promoter hypermethylation (**Fig. 3, F to H**) – also exhibited *EXO1* dependency (**Fig. 4E**), extending this finding to epigenetically driven RER deficiency. Although *EXO1* has previously been described as a *BRCA1* synthetic lethal gene (*36*), *EXO1* dependency was not observed in *BRCA1*-mutant tumor cells subjected to whole-genome CRISPR screens (**Fig. 4E**). Recent development of a small-molecule EXO1 inhibitor ART5537 and its assessment across a panel of *BRCA1*-mutant cell lines also showed that *BRCA1*:*EXO1* synthetic lethality is not a highly penetrant effect (*37*). Consistent with this, we found that Δ*RNASEH2A*, Δ*RNASEH2B*, and Δ*RNASEH2C* clones were all highly sensitive to EXO1 inhibition with ART5537 compared to wild-type cells (*P* < 0.0001), whereas Δ*BRCA1* cells showed only weak sensitivity **(Fig. 4, F and G)**. Extending this analysis to a broader panel of isogenic RPE1 HR gene knockouts confirmed that Δ*RNASEH2* clones had the greatest reduction in clonogenic survival upon EXO1 inhibition, consistent with the parallelized CRISPR screens.

Other strong dependencies that arose from the screen included the *FANCM:RNASEH2A* and *FAAP24:RNASEH2A* synthetic lethal effects, shared with Δ*BRCA1* cells only. Several lines of evidence supported these interactions. First, reanalysis of previously published CRISPR screens in two independent RPE1 Δ*FANCM* clones (*38*) revealed *RNASEH2A* as an SL dependency (CCA >1.0 and BF >8) **(fig. S8A)**. Second, siRNA co-depletion of *FANCM* or *FAAP24* with *RNASEH2A* or *RNASEH2B* reduced clonogenic survival and cellular fitness in independent HeLa and MCF10A cellular models **(fig. S8, B to D)**. Third, expression profiles of FA pathway genes in our isogenic RPE1 models showed a consistent, significant upregulation of *FANCM* and *FAAP24* in addition to other core FA pathway genes in *RNASEH2* knockout clones compared to wild-type controls **(fig. S8E)**, in keeping with the expected transcriptional buffering (*39*) of *FANCM* and *FAAP24* as a compensatory upregulatory response to *RNASEH2* defects. Lastly, chemical inhibition with the FANCM inhibitor FANCM-BTR-PPI-IN-1 reduced clonogenic survival in Δ*RNASEH2A* cells (*P* = 0.011) **(fig. S8, F and G)**. Both *EXO1* and *FANCM* are thus potential unique therapeutic targets for RER-deficient tumors, making it clinically important to distinguish this subset from some HRd tumors.

## Discussion

We identify RER deficiency as a biomarker-defined repair-deficient state in human cancer and show that mutational scars can uncover clinically actionable repair phenotypes that are not apparent from conventional gene-centric analyses. We define its mutational scars, establish *RNASEH2C* promoter hypermethylation as a recurrent mechanism of pathway inactivation, and develop a genomic classifier that identifies affected tumors across cancer types. Together, these findings extend the concept of mutational scar-based diagnostics beyond established paradigms such as HRD and MMRD, demonstrating how genome-wide mutational profiling can reveal therapeutically relevant biological states that are invisible to conventional driver-based approaches.

Replication stress is increasingly recognized as a fundamental vulnerability in cancer, yet clinically useful biomarkers of specific replication-stress states remain limited. Our findings suggest that RER deficiency represents one such state that can be identified directly from tumor genomes. The ability of RERDetect to identify tumors lacking identifiable genetic drivers highlights the value of phenotype-based diagnostics, which can capture biologically convergent repair defects regardless of their underlying mechanism.

Notably, a single repair defect can generate multiple classes of mutational scar through distinct downstream processing routes. TOP1-mediated cleavage of embedded ribonucleotides underlies the characteristic InD4c deletions, whereas the SBS3-like and structural variation components likely reflect replication-gap formation and tolerance of secondary lesions rather than bona fide HR failure (*40*). Recent studies have linked SBS3 to translesion synthesis across ssDNA gaps generated during replication stress, providing a plausible explanation for its emergence in HR-proficient RER-deficient cells (*41-43*). Together, these findings suggest that RER deficiency generates a coherent genome-wide mutational phenotype arising from persistent ribonucleotide-associated replication stress.

The identification of *FANCM/FAAP24* and *EXO1* as synthetic lethal dependencies further supports RER deficiency as a distinct biological state rather than a purely mutational classification. Both pathways function in the management of replication-associated lesions and ssDNA gaps, suggesting that RER-deficient cells rely on compensatory mechanisms to tolerate persistent ribonucleotide-induced replication stress. FANCM suppresses R-loops and stalled replication forks and has recently been implicated in limiting ssDNA gap formation, raising the possibility that its loss amplifies the replication stress generated by unrepaired ribonucleotides (*44, 45*). EXO1 similarly promotes the processing and expansion of ssDNA gaps required for ATR activation and survival of repair-deficient cells, suggesting that RER-deficient tumors may depend on EXO1-mediated gap processing to maintain viability in the presence of persistent ribonucleotide-associated lesions (*46, 47*). Although the precise mechanisms remain to be defined, these findings support a model in which RER deficiency creates a chronic replication-stress state that is buffered by FANCM- and EXO1-dependent pathways.

The discovery of *RNASEH2C* promoter hypermethylation as a novel driver of RER deficiency provides a clear example of how repair dysfunction can arise through mechanisms that evade conventional gene-centric diagnostics. It demonstrates that mutational phenotypes can reveal biologically equivalent repair-deficient states irrespective of their underlying genetic or epigenetic cause. That *RNASEH2* hypermethylation appears enriched among hypermutators, which rarely carry genetic *RNASEH2* drivers, raises the possibility that complete, sustained loss of RNase H2 activity, possibly compounded by replication stress, is required to generate extreme ribonucleotide burdens. The observation that most RERDetect-high tumors lack identifiable genetic drivers suggests that mutational scars may provide a more comprehensive measure of pathway dysfunction than genomic alterations alone.

Clinically, our findings establish RERd as a distinct and potentially actionable molecular subtype of cancer. RERDetect enables the identification of this phenotype directly from whole-genome sequencing data, capturing tumors with convergent biological states arising through diverse genetic and epigenetic mechanisms. Given the distinct dependencies of RER-deficient tumors, accurate discrimination of RERd from HRD has direct treatment implications. For example, synthetic lethal targets such as *POLQ* and *CIP2A* appear specific to HR-deficient cancers, whereas *EXO1* represents a uniquely strong vulnerability in RER deficiency. Since *RB1* co-loss, which is frequent in *RNASEH2B*-deleted tumors (*48, 49*), can induce PARPi resistance while retaining ATRi sensitivity (*25*), EXO1 inhibition may provide a potential alternative for this subset of RER-deficient cancers. Future studies will be required to determine whether RERd predicts response to PARP inhibition, ATR inhibition, or the additional synthetic lethal targets identified here.

More broadly, RERDetect provides a deployable biomarker for a replication-stress-associated repair-deficient state that is frequently missed by conventional genomic testing. These findings demonstrate how mutational scars can reveal clinically actionable biological states that are invisible to gene-centric diagnostics, extending the role of genome-wide mutational profiling beyond the identification of underlying driver events to the direct detection of therapeutically relevant tumor phenotypes.

## Acknowledgments

We gratefully acknowledge the participants of the National Genomic Research Library (NGRL), whose contributions made this research possible. Secure access to the NGRL under project ID RR239 was provided by Genomics England, which delivers the NGRL in partnership with NHS England, and is wholly owned by the UK Department of Health and Social Care. The NGRL contains participants’ health data collected by the NHS as part of their care, along with samples and data from their participation in research, for which fully informed consent has been obtained. This includes genomic and clinical data provided through the NHS Genomic Medicine Service, as well as data obtained through research studies, including the 100,000 Genomes Project and the Generation Study, both of which are delivered in partnership with the NHS, and from other research cohorts involving external collaborators.

This publication and the underlying study have been made possible partly based on data that Hartwig Medical Foundation has made available to the study through the Hartwig Medical Database.

This research was supported by the Cambridge BRC Cell phenotyping Hub.

The MASTER program is supported by the NCT Overarching Clinical Translational Trial Program, the NCT Heidelberg Molecular Precision Oncology Program, and DKTK.

## Funding

Sir Jeffrey Cheah Foundation (GCCK)

Cancer Research UK Advanced Clinician Scientist Award (C60100/A23916) (SNZ)

Dr. Josef Steiner Cancer Research Award 2019 (SNZ)

Basser Gray Prime Award 2020 (SNZ)

CRUK Pioneer Award (C60100/A23433) (SNZ)

CRUK Grand Challenge Award (C60100/A25274) (SNZ)

CRUK Early Detection Project Award (C60100/A27815) (SNZ)

National Institute for Health and Care Research (NIHR) Professorship (NIHR301607) (SNZ)

NIHR Cambridge Biomedical Research Centre (BRC-1215-20014, NIHR203312) (SNZ)

Breast Cancer Now Toby Robins Research Centre (CJL, ANJT)

Cancer Research UK Programme Grant (CJL, ANJT)

The views expressed are those of the authors and not necessarily those of the NIHR or the Department of Health and Social Care.

## Author contributions

Conceptualization: GCCK, SNZ, CJL

Investigation: PJT, SB, GCCK generated and validated isogenic knockouts, performed mutation accumulation and sequencing experiments with help from SJZ.

Analysis: PJT analyzed the sequencing data with guidance and input from GCCK, DB, SJZ, GR, AD, YM, HRD.

CRISPR screen and validation: RB, FFS, CJL designed and performed the CRISPR screen. PJT validated screen targets with help from SJZ and RB.

Visualization: PJT, GCCK, SB, SJZ, RB, FFS, JC, YM, HRD, CJL

Funding acquisition: GCCK, SNZ, CJL, ANJT, SJP Supervision: GCCK, SNZ

Writing – original draft: PJT, GCCK, SNZ, CJL Writing – review & editing: All authors

## Competing interests

GCCK, GR, AD, HRD, and SNZ are inventors on patent applications encompassing the code and intellectual principle of the HRDetect algorithm (PCT/EP2017/060294) (HRD, SNZ), clinical use of signatures (PCT/EP2017/060289) (HRD, SNZ), clinical predictors (PCT/EP2017/060298) (HRD, SNZ), rearrangement signature methods (PCT/EP2017/060279) (SNZ), mutational signatures method (PCT/EP2023/056078) (AD, SNZ), indel signature methods (PCT/EP2024/077959) (GCCK, SNZ) and PRRDetect algorithm (PCT/EP2024/078030) (GCCK, GR, SNZ). CJL receives and/or has received research funding from AstraZeneca, Merck KGaA, and Artios; received consultancy, SAB membership, or honoraria payments from Syncona, Sun Pharma, Gerson Lehrman Group, Merck KGaA, Vertex, AstraZeneca, Tango, 3rd Rock, Ono Pharma, Artios, Abingworth, Tessellate, Dark Blue Therapeutics, Pontifax, Astex, Neophore, GlaxoSmithKline, Dawn Bioventures, Blacksmith Medicines, and FoRx; has stock in Tango, Ovibio, Hysplex, Tessellate, and Ariceum. He is also a named inventor on patents describing the use of DNA-repair inhibitors, including ATR inhibitors, and stands to gain from their development and use as part of the ICR ‘Rewards to Inventors’ scheme and also reports benefits from this scheme associated with patents for PARP inhibitors, paid into CJL’s personal account and research accounts at the Institute of Cancer Research. ANJT reports personal honoraria for advisory boards from AstraZeneca, ForEx Therapeutics, Nauge Therapeutics and Artera, and honoraria for advisory boards to either The Institute of Cancer Research or King’s College research accounts from AstraZeneca Page Therapeutics, Ellipses, Tango Therapeutics, Guardant Health, Merck KGaA, as well as speaker fees from AstraZeneca and Permanyer Mexico. ANJT reports financial support for research from AstraZeneca, Myriad Genetics, Merck KGaA and compensation for travel expenses from AstraZeneca. ANJT reports benefits from ICR’s Rewards for Discoverers Scheme associated with patents for PARP inhibitors in *BRCA1*/*2* associated cancers, paid into research accounts at The Institute of Cancer Research and to ANJT’s personal account. SJP makes the following disclosures: SJP is a named inventor on patents describing the use of DNA repair inhibitors and treatment of cancers with reversion mutations and stands to gain from their development and use as part of the ICR “Rewards for Discoverers” scheme. SAB membership: VUS Genetics. All other authors declare no competing interests.

## Data, code, and materials availability

Raw sequence files from the hTERT-RPE1 mutation accumulation experiment are deposited at the European Genome-Phenome Archive with accession number: EGADXXXXXXXX. Mutation calls have been deposited at Mendeley: doi:xxxxxxx. RPE1 cells can be obtained directly from the authors. The curated data are available for general browsing from our reference mutational signatures website, Signal (https://signal.mutationalsignatures.com). All other data were previously published (*50, 51*).

The results of RPE1 experimental signatures can be browsed at https://signal.mutationalsignatures.com/explore/experimental?study=7.

Data from the National Genomic Research Library (NGRL) used in this research are available within the secure Genomics England Research Environment. Access to NGRL data is restricted to adhere to consent requirements and protect participant privacy (*52*). Data used in this research include: WGS and substitution, indel, and rearrangement mutational profiles of *n*=12,007 samples. Details on how these datasets were derived are described in Methods. Access to NGRL data is provided to approved researchers who are members of the Genomics England Research Network, subject to institutional access agreements and research project approval under participant-led governance. For more information on data access, visit: https://www.genomicsengland.co.uk/research.

The R source code of RERDetect can be obtained from https://github.com/Nik-Zainal-Group/RERDetect.

