## Supplementary Materials for "Global signatures of Ribonucleotide Excision Repair deficiency and its therapeutic vulnerabilities"

Cell line models

A summary of all human cell line models used in this study is provided in **Table S1**. RPE1 and MCF7 cells were grown in DMEM/F12 medium (Thermo Fisher Scientific, 11320033) supplemented with 10% FBS. hTERT-RPE1 *ΔTP53* cell (a gift from M. Tarsounas, Department of Oncology, University of Oxford, UK) was generated from a previous study (*13*). HAP1 cells were grown in Iscove’s Modified Dulbecco’s Medium (Gibco^TM^, 12440053) containing L-Glutamine and 25 mM HEPES, supplemented with 10% FBS. MCF10A cells were grown in DMEM/F12 supplemented with 5% horse serum, EGF, hydrocortisone, cholera toxin, and insulin. HeLa cells were grown in DMEM, high glucose, GlutaMax supplement, pyruvate (Gibco^TM^, 11960044) supplemented with 10% FBS. HeLa cell lines (gifted from A. Jackson, Institute of Genetics and Cancer, University of Edinburgh) were generated from a previous study (*13, 53*). HCC202 is an epithelial cell line isolated from a mammary gland invasive ductal carcinoma of TNM stage IIIA, grade 3. HCC202 was grown in RPMI 1640 medium (Gibco^TM^, 11875093) supplemented with 10% FBS. All cells were grown at 37°C and 5% CO_2_ in a humidified incubator.

To generate the remaining isogenic CRISPR-edited cell lines in hTERT-RPE1 *ΔTP53*, 200k cells per each edit were electroporated with pre-formed ribonucleoprotein complex (RNP with final concentrations of 120 *p*mol gRNAs and 100 *p*mol Alt-R Cas9) in supplemented SE buffer, using nucleocuvette and AMAXA 4D-Nucleofector (Lonza, V4XC-1032) on program EH-158 according to manufacturer’s instructions. Following electroporation, cells were incubated for 5 min at room temperature and 10 min at 37°C in 80 μl Opti-MEM™ I Reduced Serum Medium (Gibco™, 31985062). They were then re-plated into fully supplemented DMEM/F12 medium to recover for 48h before limiting dilution on 96-well plates to isolate single-cell clones. Targeted clones were screened for successful frameshift edits for knockouts via Sanger sequencing. All gRNAs, sequencing primers, and genotypes of the cell line models used in this study are summarized in **Tables S1 and S2**.

Cell-cycle analysis

Cell cycle analysis was performed using the Propidium Iodide Flow Cytometry Kit (Abcam, ab139418) according to the manufacturer’s protocol. Cells were rinsed with PBS, trypsinized, and quenched with pooled culture media to generate a single-cell suspension. Cells were centrifuged at 500 x g for 5 min. The supernatant was discarded, and the pellet was resuspended in 1 mL PBS. Cells were fixed using 66% ethanol on ice. At room temperature, cells were centrifuged at 500 x g for 5 min. Supernatants were discarded, and pellets were washed again with 1 mL PBS. Cells were collected and resuspended in 200 µL Propidium Iodide + RNase Staining Solution. Samples were incubated at 37°C in the dark for 30 min then placed on ice for flow cytometry analysis.

Samples were analyzed using an acoustic flow cytometer Attune NxT at the BRC Cell Phenotyping Hub. Unstained control cells were used to determine gating of single cells using forward and side scatter. Fluorescence on cells was excited by the blue laser (488 nm) and detected using BL2 with a 695/40 band-pass filter. The percentage of cells in each cell cycle phase was analyzed using the Watson cell-cycle model on FlowJo (v.10.10.0). Three independent biological replicates were used for statistical analysis conducted on GraphPad prism (v.10.4.1).

Dose-response assays

Cells were seeded into 96-well plates at a density of 300 cells per well in 50 µL of complete medium. Cells were allowed to adhere for 24 h in the incubator before treatment. The following drug treatments with a range of doses along with DMSO vehicle control were prepared: ATR inhibitor (AZD6748; Cayman Chemical 21035) 0 to 4 μM, PARP inhibitors Olaparib (APExBio, A4154) 0 to10 μM and Talazoparib (Selleck Chemicals, S7048) 0 to 500 nM, and TOP1 inhibitor Camptothecin (MedChemExpress, HY-16560) 0 to 10 nM. Drugs were added to the appropriate wells in 50 μL medium. CellTiter-Glo Luminescent Cell Viability Assay (Promega, G7570) was conducted after 7 days when control wells reached 95% confluence. Briefly, 100 μL of the assay reagent was added to the wells, and the plates were gently agitated to ensure thorough mixing. The luminescence signal was then measured using a PHERAstar *FS* microplate reader (BMG Labtech) according to the manufacturer’s guidelines. Signals were blanked and normalized to respective solvent controls. Statistical analysis was conducted using GraphPad prism (v.10.4.1).

Immunoblotting

Proteins were extracted from cell pellets using RIPA buffer (Sigma-Aldrich, R0278) supplemented with protease and phosphatase inhibitors (Thermo Scientific, 87786) and 0.1% benzonase (Merck, E1014) for 15 min on ice. Lysates were then centrifuged at 15,871 x g for 10 min at 4°C. Equal amounts of protein from whole-cell extracts (WCE) were separated by SDS–PAGE on 4–12% NuPAGE gels and transferred to PVDF membranes. Membranes were probed in 5% milk (w/v; nonfat dried milk powder), PBS + 0.1% Tween-20 (v/v) with the following antibodies: mouse anti-RNase H2A (Proteintech, 67967-1-Ig, 1:1,000); rabbit anti-RNase H2B (Proteintech, 84097-6-RR, 1:1,000); rabbit anti-RNase H2C (Proteintech, 16518-1-AP, 1:1,000), and rabbit anti-HSP90 (Proteintech, 13171-1-AP, 1:2,000). For detection, we used goat anti-mouse immunoglobulins/HRP (Abcam, ab150114 1:2,000); anti-rabbit IgG, HRP-linked antibodies (Abcam, ab205718, 1:2,000); Clarity Western ECL Substrate (Bio-Rad, 1705060) and an iBright FL1500 Imaging System (Thermo Fisher).

Alkaline gel electrophoresis

To determine the presence of excess genome-embedded ribonucleotides in nuclear DNA, alkaline gel electrophoresis of RNase H2-treated genomic DNA was conducted with modifications to previous protocols (*53*). In brief, DNA was extracted using the GenElute-E Single Spin Cell Culture kit (Sigma Aldrich, EC400). 1 µg of DNA was incubated with 15 U of purified recombinant human RNase H2 (NEB, M0288) and 0.25 µg of DNase-free RNase (Roche, 11119915001) for 1 h at 37 °C in 100 µl ThermoPol Reaction Buffer (NEB, B9004). Samples were cleaned up using DNA Clean & Concentrator-5 kit (Zymo Research, D4014) according to protocol. Samples were run on a 0.8% agarose gel in 50 mM NaOH, 1 mM EDTA in the dark. After overnight electrophoresis, the gel was neutralized in 0.7 M Tris–HCl pH 8.0, 1.5 M NaCl for 30 min and stained with SYBR Gold (Invitrogen, S11494) diluted 1:10,000 in TAE buffer for 10 min. Imaging was performed with Bio-Rad Gel Doc XR+ System and densitometry analyzed using RStudio (v.2024.04.1+748).

RNase H2 activity assays

RNase H2 activity assays were adapted from that described previously (*54*). Whole-cell lysate was used. Washed cell pellets were resuspended in lysis buffer (50 mM Tris HCl pH 8, 280 mM NaCl, 0.5% NP-40, 0.2 mM EDTA, 0.2 mM EGTA, 10% glycerol vol/vol, 1 mM DTT, and 1mM PMSF). Cells were incubated on ice for 10 min. Addition of an equal volume of dilution buffer (20 mM HEPES pH 7.9, 10 mM KCl, 1 mM EDTA, 10% glycerol, 1 mM DTT, 1 mM PMSF) was followed by another incubation on ice for 10 min. Cell debris was pelleted by centrifugation at 17,000 x g for 10 min at 4 °C. Supernatant was kept and quantified using the Pierce BCA protein assay kit (Thermo Scientific, 23225).

For the activity assay, 10 μM of oligonucleotides (GenScript) were annealed by heating for 5 min at 95 °C then gradually cooling to room temperature 18 °C in 60 mM KCl, 50 mM Tris-HCl pH8. 100 ng/µL whole-cell lysate was dispensed into a 96-well plate with a total of 100 µL per well in a buffer of 60 mM KCl, 50 mM Tris HCl pH8, 10 mM MgCl2, 0.01% BSA, 0.01% Triton-X100. 250 nM of corresponding substrate was added to each well. For each condition, triplicate reactions were measured using PheraStar *FS* (BMG Labtech) at 24 ± 2 °C with shaking at 60 rpm every few minutes over 120 min. Fluorescence was read using a 480 nm excitation filter and 535 emission filter. All conditions were subtracted by its associated background substrate fluorescence with no whole-cell lysate and normalized to the positive control substrate with expected maximum fluorescence.

Homologous Recombination (HR) activity assays

The extrachromosomal reporter of long HR activity assay was conducted as previously described (*11*). The long template HR (pCMV_LHR) plasmid was used to measure HR activity. Plasmids were delivered into RPE1 cell lines using electroporation with AMAXA 4D-Nucleofector (Lonza, V4XC-1032) on program EN-138. Plasmid to firefly ratio was calculated as 1:3.5 according to mass in total of 400 ng per electroporation reaction. 225,000 cells were used per reaction, and 75,000 cells were seeded per well in a 96-well plate. 6 technical replicates were used. The assay was terminated 24 h and 30 min post-electroporation. Nano-Glo Dual Luciferase Reporter Assay System (Promega, N1620) was used to quantify the NanoLuc luminescence signals using PheraStar *FS* (BMG Labtech). Luminescence signals were normalized to values from respective firefly transfection control and to wild-type cell lines.

Methylation quantification

Extracted DNA from HCC202 and MCF7 was bisulfite-converted using the EZ DNA Methylation-Lightning Kit (Zymo Research, D5030). Converted DNA was then amplified by PCR with methylation-specific primers targeting the *RNASEH2C* promoter (**Table S15**). PCR products were sequenced using Sanger Sequencing. QC of sequencing results included sequence trimming to a minimum Phred quality score of 20, a minimum length of 30-bp, a minimum ratio of primary-peak 0.7, and a minimum percentage of non-mixed position of 40%. Sequences were aligned with a minimum length of 30-bp and a minimum bisulfite-conversion rate of 0.9. QC, alignment, and methylation quantification analysis were performed using Analysis of Bisulfite Sequencing PCR on R (ABSP, v1.2.2) (*55*).

Genome wide CRISPR-Cas9 screen and analysis

Cas9-expressing RPE1 *ΔTP53* isogenic knockout cell lines were transduced with the human genome-wide lentiviral CRISPR nuclease (CRISPRn) library (Addgene, 67989) (*56*). Cells were infected at a multiplicity of infection (MOI) of 0.3 (to avoid multiple sgRNA integrations per cell) and a minimum of 500x representation per sgRNA was maintained throughout the experiment. Following selection with 20 µg/ml puromycin for 72 h, cells were seeded into DMSO-containing media, and a T0 sample was collected. Cells were passaged in DMSO-containing media for 14 days, after which the final T1 sample was collected. Genomic DNA was extracted from the T0 and T1 samples, and sgRNA were amplified using 25 cycles of PCR with Illumina primers before targeted sequencing to a target depth of 500X on NovaSeq 6000 (Illumina) to generate count data. For synthetic lethal analysis, BAGEL2 (*57*) and CRISPR Count Analysis (*33*) (CCA) (<https://github.com/tohsumi-repare/cca>) were performed on each DMSO screen sample. Genes with a BAGEL2 Bayes Factor (BF) >0 and CCA score >0.5 were considered to show mutant-selective dependency effect (*57*).

Clonogenic assays and siRNA transfection

TriFECTaDicer-substrate small interfering RNA (DsiRNA) duplexes targeting *RNASEH2A*, *RNASEH2B*, *FANCM*, and *FAAP24* (IDT) were transfected into HeLa and MCF10A (**Table S16**). HeLa cells were transfected using Lipofectamine RNAiMAX according to the reverse transfection protocol (Thermo Fisher Scientific, 13778150). MCF10A cell lines were electroporated with 3 μM of DsiRNA using AMAXA 4D-Nucleofector (Lonza, V4XC-1032) on program EN-138. Transfected cells were allowed to recover for 24 h. Knockdown efficiency was verified by qPCR using the SYBR Green Fast Advanced Cells-to-CT kit (Thermo Fisher, A35379). cDNA amplification and relative quantification was run on QuantStudio 3 Real-Time PCR Systems (Applied Biosystems). qPCR primers used are denoted in **Table S15**. Relative gene expression levels were analyzed using the ΔΔCt method. Additional validation of sufficient gene target knockdown was verified using immunoblotting. Cell pellets for immunoblotting were collected 4-5 days post-electroporation.

For drug treatments, cells were seeded into triplicate wells and treated after 24 h with EXO1i ART5337 (*37*) (5 μM) or FANCM inhibitor FANCM-BTR-PPI-IN-1 (*58*) (25 μM) for 6 days before crystal violet staining.

For the clonogenic assay, 500 cells were seeded into triplicate wells after 24 h in 6-well plates. Cells were grown for 7-10 days, washed with PBS, and fixed with 100% methanol. After drying, cells were stained with 0.5% crystal violet in 25% methanol for 30 min. Colonies were imaged using iBright FL1500 Imaging System (Thermo Fisher) in Universal mode and counted using the associated analysis software.

Mutation accumulation experiment and sequencing

All edited clones were expanded and maintained in culture for over 100 days (approximately 80-100 cell doublings) to allow for mutation accumulation in the edited genetic backgrounds. A second round of single cell limiting dilution was then performed to isolate 2-3 daughter subclones per edit genotype, providing a bottleneck to capture mutations that had occurred since the isolation of the initial edited parental clones.

Genomic DNA was isolated from all samples using *Quick*-DNA Miniprep Plus Kit (ZymoResearch, D3024) following the manufacturer's protocol. WGS libraries were prepared and sequenced with a paired-end 150-bp configuration on Illumina NovaSeq 6000 platform by Novogene (Beijing, China), aiming for an average genome-wide sequencing depth of 25X per sample.

Somatic variant calling

Short reads were aligned to GRCh38/hg38 using BWA-MEM 0.7.17-r1188. Substitutions and indels in genetic reference clones and daughter subclones were called as previously described (*59*). Post-processing filters were applied to improve the specificity of mutation-calling. Specifically, for SNV calls by CaVEMan (*60*) (v1.13.15), we used CLPM == 0 and ASMD ≥ 140. To reduce false positive calls by Pindel (*61*) (v3.2.0), we used QUAL ≥ 250 and REP < 10. Variant allele fraction (VAF) distribution for each subclone was examined (*i.e*., estimation of mutation density and mutation burden). *De novo* substitutions and indels in subclones were obtained by subtracting from respective parental clones whenever available, or by removing mutations shared among subclones. *De novo* mutation counts are provided in **Table S3**. Variant allele fraction (VAF) distributions for each subclone were examined, applying VAF≥ 0.2 for substitutions and VAF ≥ 0.25 for indels to exclude subclonal variants. No VAF filter was applied for rearrangements.

Mutational signature analysis of experimental samples

The derivation of gene edit-associated mutational signatures with cosine similarity, bootstrapping and background subtraction was performed using published framework (<https://github.com/xqzou/COMSIG_KO>) (*50*). In short, we first (i) determined the background mutational signature in background control by aggregating the unedited and untreated subclone mutational profiles; (ii) determined the difference between the mutational profiles of the edited clones and background mutation profile; and if an edit generates a signature, we (iii) removed background mutation profile from the mutation profile of the edited subclone (**Table S1**).

Experimentally derived signatures were compared to published reference signatures (*62, 63*) using signature.tools.lib from <https://rdrr.io/github/Nik-Zainal-Group/signature.tools.lib/>. Indel signatures were analyzed using <https://github.com/Nik-Zainal-Group/indelsig.tools.lib/>.

Global transcriptomics

Total RNA was extracted using Direct-zol RNA Miniprep Plus (Zymo Research, R2073) according to the manufacturer’s protocol. cDNA libraries were prepared and sequenced by Novogene (Beijing, China) on Illumina NovaSeq 6000 platform. At least 20 million 150-bp paired-end reads were generated for each sample. Spliced Transcripts Alignment to a Reference (STAR, v.2.5.0.a) (*64*) was used to map RNA-seq data to the reference genome GRCh38. Reads overlapping gene features were counted using featureCounts (v.2.0.1) (*65*). Raw count matrices were analyzed using the DESeq2 R package (*66*). Counts were normalized by median-of-ratios method and underwent variance stabilizing transformation (VST). Genes of <10 reads in total were removed. Gene ontology analysis was performed using R package clusterProfiler (v.4.12.6) (*67*). SLM^HRD^ scores were computed from DESeq batch normalized RNA-seq expression values. The 73-gene SLM synthetic lethal genes were obtained from the original publication (*39*). GSVA (v.1.52.3) was applied to the expression matrix to obtain an enrichment score.

Datasets

We analyzed samples from three large pan-cancer whole-genome cohorts including the Genomics England (GEL) 100kGP data release v17 (GEL, <https://www.genomicsengland.co.uk/>), the ICGC cohort, and the Hartwig Medical Foundation metastatic cohort samples as previously described (*63*).

Other datasets include the SCAN-B cohort of 254 triple-negative breast cancers (*26*) and a subset of the MASTER cohort of 1,876 patients, most of whom have rare cancers and soft tissue cancers as described previously (*22*).

Substitutions were classified according to their trinucleotide context into 96-channel catalogues using signature.tools.lib (v.2.4.6). Indels were categorized into 89-channel indel context using indelsig.tools.lib (v1) (*10*), and structural variants were categorized into whether they occurred in a cluster of other SVs or not, by their size and type into 32-channel catalogues (*2*).

Indel signatures were extracted and fitted using FitMS in signature.tools.lib from <https://github.com/Nik-Zainal-Group/signature.tools.lib/> with 500 bootstraps, maximum 2 rare signatures per sample, and a gini threshold using the new 89-channel classification system in 12,007 samples in GEL (*10*). Substitution signature exposure data was obtained from a previously published paper for all samples that also had available indel signature exposure data (*63*).

HRd algorithms and classifiers

HRDetect (*20*) probability scores were obtained from a previously published paper (*63, 68*). The algorithm was run on default parameters. Samples with HRDetect probabilities ≥0.7 were classified as high, scores <0.7 and >0.1 as intermediate, and ≤0.1 as low. Signature Multivariate Analysis (SigMA) (*19*) was performed using the SigMA R package (<https://github.com/parklab/SigMA>, v1.6). The Classifier of Homologous Recombination Deficiency (CHORD) (*18*) was implemented using the CHORD R package (<https://github.com/UMCUGenetics/CHORD>, v2.3). A default threshold of ≥0.5 was applied to identify HRd samples. HRProfiler was implemented using the R package (<https://github.com/AlexandrovLab/HRProfiler>, v.0.0.2) using the default parameters. A default threshold of ≥0.5 was applied to identify HRd samples. SigMA, CHORD, and HRProfiler were run on 60 GEL breast samples that had available HRDetect scores.

Sequence logo

Genomic sequences 10-bp upstream and downstream from the position of each 2-bp deletions were obtained and classified from the Pindel VCF files. Deletions from the Pindel VCF files were left aligned. Genomic sequences were reverse complemented when the deleted dinucleotide contained an adenosine (A), except for AT or TA. Example (2-bp deletion underlined, trinucleotide motif in bold):

(1) 5’ TACCCCCAAAT**TCT**GTTGGGT3’ left-aligned 2-bp deletion at microhomology
(2) 5’ GCATTTTAAAT**ACA**GTCATTT3’ left-aligned 2-bp deletion at microhomology
(3) 5’ AAATGAC**TGT**ATTTAAAATGC3’ right-aligned 2-bp deletion at microhomology reverse complemented sequence in (2).

All deletions were then right aligned so that the 2-bp deletion started at position 0. Sequences were converted to bit score matrices and logos were generated using Logomaker (v.0.8) (*69*). Two-base-pair deletions were randomly simulated across the mappable genome excluding blacklisted regions. The enrichment of 2-bp deletions at TNT motifs and transcribed regions compared to randomly simulated frequencies was calculated using Fisher’s exact test.

Indel hotspots

Indels from 48 cancers with InD4c exposure of at least 150 and proportional InD4c exposure at least 0.2 were binned into 500kb windows across the entire genome. Only indels from the three main InD4c indel channels (Del(2,5):M1, Del(2,8):U(1,2):R(2,4), and Del(2,4):R1) were used. In total, 26,609 indels were found across 5,620 windows. Windows with at least 10 indels were defined as indel hotspots. Background indels of similar genomic context was randomly generated using SigProfilerSimulator (v.1.2.2) (*70*) with default parameters and 20 repeats per sample. One sample t-test was used to determine windows with significantly higher numbers of indels than expected. Correlations between indel hotspots with R1 hotspots (*71*), R loops (*72*), origins of replication (*73*), and BrdU peaks (*74*) was conducted using topography.tools.lib (<https://github.com/Nik-Zainal-Group/topography.tools.lib>).

RERDetect

*Training set*

We trained RERDetect, a lasso logistic regression model on 227 cell lines consisting of 19 homozygous *RNASEH2* knockout samples and 208 non-*RNASEH2* mutated samples as ground truth. All cell line genotypes were confirmed by WGS. Ground truth negative samples included *BRCA1*, *BRCA2* knockout cell lines, MMRd cell lines, chronic drug treated cell lines and control wild-type cell lines. Ground truth negative samples do not have mutations in *RNASEH2* and were included to increase variation in genomic features.

*Feature engineering*

To create the classifier, we explored various combinations of input features. These were divided into mainly mutation contexts and signature exposures. Mutation contexts included three main somatic mutation categories (SNV, indels, and structural variants). The mutation contexts used as inputs included (1) Proportion of mutational catalogues with 96 substitution channels, 476 indel channels, and 16 rearrangement channels and (2) Proportion of mutational catalogues with 96 substitution channels, 89 indel channels, and 16 rearrangement channels. The 96 substitution channels include the SNVs of C>A, C>G, C>T, T>A, T>C, or T>G with flanking 5’ and 3’ nucleotides. The 89 indel channels include 1-bp C, 1-bp T, >=2-bp insertions and deletions, deletions at microhomology, and complex indels. The 476 indel channels include more comprehensive subcategories of the 89 indel channels with higher resolution. The 16 rearrangement channels include deletions, tandem-duplication, and inversions, at 1-10 kb, 10-100 kb, 100 kb-1 Mb, 1-10 Mb, and >10Mb, and translocations.

Input features for signature exposures included substitution, indel, and rearrangement signatures using (3) absolute values and (4) proportions. To obtain signature exposures for experimental samples, the mutational catalogues were fitted with 200 bootstraps to the experimentally derived mutational signatures associated with HR-deficiency, MMR deficiency, and background wild-type signatures. Input features that involved proportions were determined by dividing by the total substitution, indel and rearrangement burden depending on the mutation type. All features were log2 and z-score normalized using the formula x’ = (x - μ) / σ. Feature selection was conducted using a Wilcoxon test using a *P*-value cutoff of < 0.01 to minimize the number of input features.

*Training procedure*

For all models, the selected features were trained using the lasso logistic regression model from the glmnet package (v.4.1.10) in R. A total of 10 training iterations was conducted to determine feature stability. For each iteration, the cohort was partitioned randomly and uniquely to 70% training and 30% testing with maintained proportions of positive and negative samples in each iteration (**Table S8**). Within each iteration, a 10-fold cross validation was conducted as part of the cv.glmnet function for tuning hyperparameter *λ* defining the strength of the penalty using log loss as a performance metric.

Models were then retrained and tested on all 227 samples for a final whole model with a 10-fold cross validation for the final coefficients. Several performance metrics were gathered for each model including training accuracy of the final whole model, median accuracy on the test set, standard deviation of *λ,* median coefficient standard deviation, and median AUC of the test set and final whole model (**Table S9**).

*Performance validation*

All models were then validated on a cancer cohort of 549 samples with 528 control cancers and 21 unique cancers with *RNASEH2B* driver mutations manually verified using bam files. Negative control samples had to meet several criteria including (1) no FFPE samples, (2) No mutation in *RNASEH2A*, *RNASEH2B*, or *RNASEH2C*, (3) Gene expression of *RNASEH2* genes had to be above the mean gene expression (log2 normalized TPM counts from RNA-seq) of the entire cohort of cancers (4) minimum burden of substitutions > 250, indels > 150, and rearrangements > 25. Signature exposures were obtained from signature fitting of reference substitution, indel, and rearrangement signatures with 500 bootstraps. The AUROC, AUPRC, and F1 values were calculated for each model including models using mutation contexts and signature exposures as input features (**Table S11**). As the model using signature exposures as input features was trained on cell line experimental samples, the substitution, indel, and rearrangement signature exposures used to calculate the probability of cancer samples were SBS3, InD4c, and the average of rearrangement signatures R1, R22, R7, and R25. These cancer-derived signatures had the highest cosine similarity to the experimental derived signatures.

The best model was the one that had the best performance statistics in both the cross-validation and the independent cancer cohort. The final model consisted of 7 features, 2-bp deletions with single-nucleotide microhomology (Del2:M1; weight 3.26), 2-bp deletions occurring at short tandem repeats (STRs) of three (Del2:U2:R3), two (Del2:U2:R2), and four units (Del2:U2:R4) (weights 0.999, 0.721, and 0.412, respectively), T>A transversions in 5′-CTA-3′ (0.504) and 5′-CTG-3′ (0.101) contexts, and 10–100 kb tandem duplications (0.066). The cutoff threshold 0.3 was the threshold at which the F1 score was maximized in the validation cohort.

RERDetect in cancer cohorts

RERDetect was applied to three main pan-cancer cohorts including 7,507 GEL samples, 1,491 ICGC samples, and 2,702 metastatic Hartwig samples (**Table S12**). These samples were derived from previous datasets using minimum burdens of substitutions >250, indels > 150, and rearrangements > 25. All samples were log2 normalized and scaled by the mean and standard deviation of the independent validation cohort of 549 cancers. Drivers in *RNASEH2A*, *RNASEH2B*, and *RNASEH2C* were sought as described previously (*75*).

Identification of driver events in cancer samples

*Loss-of-function* driver mutations in *RNASEH2A*, *RNASEH2B*, and *RNASEH2C*, helicases *ATRX*, *SETX* and classical tumor suppressor genes *RB1* and *TP53* were sought in all RERDetect-high cancer samples as previously described (*75*). In brief, somatic and germline deep deletion events (homozygous deletions), structural variant events, and *loss-of-function* coding mutations resulting in a stop-gain, frameshift, splice donor, or acceptor variant, stop-lost, start-lost, and splice donor 5^th^ base variant or missense mutations were considered as potential driver events. Any homozygous deletions in *RNASEH2* were considered when the segment of the total copy number was 0. To avoid calling artefacts, the deletions were confirmed manually on IGV, and the copy number segment had to be <=4Mb long.

Evidence that both copies of the gene were inactivated in the tumor was required to call high-confidence biallelic driver mutations. This could include Loss of Heterozygosity (LOH) of the wild-type allele (estimated from copy number solutions, tumor purity, and the VAF of the mutation in the tumor), or a second mutational hit. When seeking driver events in *TP53*, in keeping with the rules for typical tumor suppressor gene drivers, inactivating mutations were only considered high confidence if there was evidence that all copies of the gene had been disrupted. However, due to the dual role of *TP53* as both oncogene and tumor suppressor gene, monoallelic missense mutations were considered as high confidence.

Public cancer dependency data

CRISPR-Cas9 gene dependency (Chronos) scores were obtained from DepMap portal (BROAD, release 24Q2, <https://depmap.org/>). Mutational catalogues were created from processed and filtered variants calls from 339 whole-genome sequenced cell lines were obtained from published article (*27*). RERDetect was run on the mutational catalogues to identify RER-deficient cell lines.

Statistics and reproducibility

All comparisons were between biologically independent samples. No data were excluded from the analyses. The experiments were not randomized. The investigators were not blinded to allocation during experiments and outcome assessment.
